# Redating of the To’aga site (Ofu Island, Manu’a) and a revised chronology for the Lapita to Polynesian Plainware transition in Tonga and Samoa

**DOI:** 10.1101/532648

**Authors:** Fiona Petchey, Patrick V. Kirch

## Abstract

Radiocarbon dating Pacific archaeological sites is fraught with difficulties. Often situated in coastal beach ridges or sand dunes, sites exhibit horizontal and vertical disturbances, while datable materials such as wood charcoal are typically highly degraded, or derived from old trees or drift wood and bone collagen rarely survives in the tropical conditions. Shell, therefore, is the most logical material for dating Pacific sites since it is resistant to alteration, can be sampled to ensure only the last few seasons of growth are represented and is often closely tied to human economic activities. However, shell radiocarbon (^14^C) dating has been plagued by interpretive problems largely due to our limited knowledge of the ^14^C cycle in near shore marine and estuarine environments. Consequently, shell dates are typically ignored in regional chronometric evaluations and often avoided for dating altogether. Recent advances in our understanding of the source of shell ^14^C content as well as the development of the first South Pacific Gyre model of changing marine ^14^C over time, combined with Bayesian statistical modelling, have now provided us with insight into the value of these shell radiocarbon dates, enabling a revision of the age of the To’aga site on Ofu Island, an early occupation site associated with the initial Polynesian Plainware period in Samoa, the earliest use of which is now dated to between 2782 and 2667 cal BP.

## Introduction

The Lapita cultural complex, of which dentate-stamped pottery is a defining component, is argued to have begun in the Bismarck Archipelago around 3350 years ago. From there it spread into previously uninhabited areas of Remote Oceania (Fig 1), reaching its eastern-most extent in the Tongan and Samoan archipelagos around 2900-2750 BP [1][2][3]. The timing of first landfall on each island group by these Lapita explorers has been the subject of a number of chronological evaluations (e.g., [4][5][6][7][8][9][10][11]) but several parts of this story remain controversial. Of note are debates over the timing of the transition from dentate-stamped Lapita ceramics to undecorated ceramics (termed Polynesian Plainware [PPW]) - a significant change in material culture that is considered to mark the onset of Ancestral Polynesian society [12].

**Fig 1:**
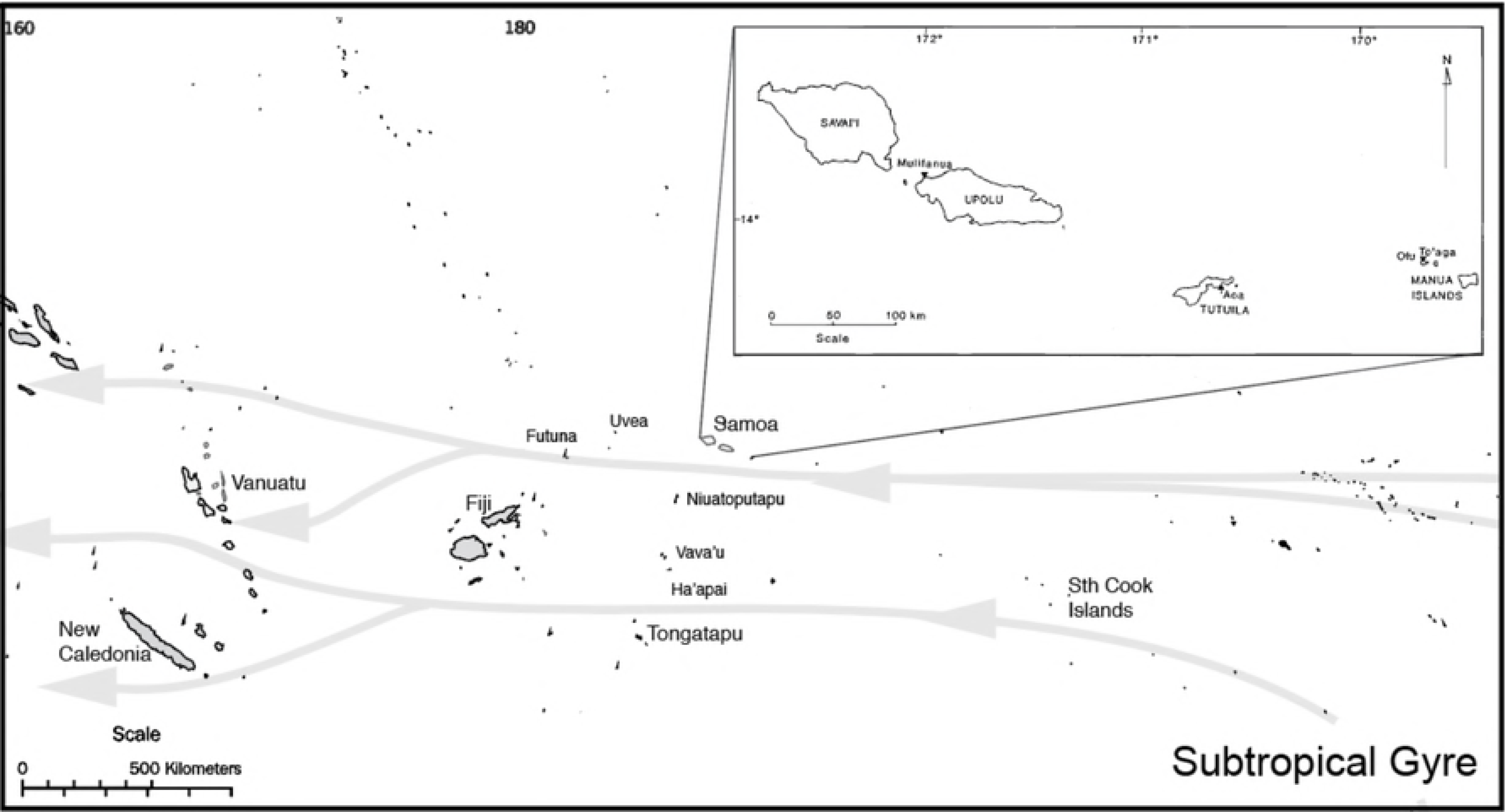
Map of the central Subtropical Gyre waters and associated island groups. Insert: Samoan Archipelago showing sites mentioned in text.

There is no doubt that Lapita colonizers reached the island of ’Upolu in Western Samoa, where the submerged coastal site of Mulifanua (SU-17-1) contains classic dentate-stamped Lapita ceramics of local manufacture [13][14]. The exact date of this settlement remains somewhat problematic because the material was obtained by dredging during the construction of a ferry terminal [15]. Radiocarbon dates (here reported at 68% probability) obtained on material collected from the dredging spoils produced a date on turtle bone and shell of 2880-2750 cal BP [16]. Comparably early charcoal dates have also been obtained from the sites of ‘Aoa (AS-21-5) on Tutuila Island, and To‘aga (AS-13-1) on Ofu Island in the Manu’a group. Dates from these sites suggested that occupation either pre-dated, or was contemporaneous with, Mulifanua [17][18][19], but neither ’Aoa or To’aga produced ceramics with dentate stamped decoration. Instead, red- and orange-slipped thin, fine tempered plainware was recovered from the deepest units of both sites. These ceramics were considered to represent a distinct marker horizon between the earliest layers and those that contained thicker, coarse-tempered pottery which became dominant after 2400 cal BP ([20] pg 91).

Rieth and Hunt [21] evaluated all available ^14^C dates from the Samoan archipelago. Their analysis “challenged the validity of the earliest dates from ‘Aoa and To‘aga” because of their low precision (standard errors in excess of ±100 years). This left only shell dates from To’aga for consideration, suggesting a ~300 year separation between the ceramics from Mulifanua and the earliest occupations at To‘aga (2500-2400 cal BP), and two other early sites at Utumea (Tutuila Island; 2500-2100 cal BP; AS-22-44) and Jane’s Camp (Upolu; 2300-2000 cal BP; SU-18-1, SU-F1-1). Addison and Morrison ([22] pg 363) subsequently concluded that the limitations of current radiocarbon technology and calibration methods, combined with the problems of stratification and mixing of sandy Pacific coastal sites, meant that further refinement of the absolute chronology for early Samoa was unlikely.

This post-2500 BP “re-colonization” date was questioned by Clark et al. [23] who presented new data from three PPW sites on the island of Ofu: Va’oto (AS-13-13), Coconut Grove (AS-13-37) and Ofu Village (AS-13-41). Combining these results in a single-phase Bayesian model populated by a combination of short-lived *Cocos nucifera* endocarp charcoal and highly precise U/Th coral dates, Clark et al ([23] pg 272) concluded that initial settlement of Ofu Island occurred between 2717 and 2663 cal BP. This date overlapped with modelled dates for the end of Lapita from sites on Tongatapu, Ha’apai, and Vava’u (2703-2683 cal BP) where a chronological progression from Lapita to PPW sites had been identified [24], suggesting settlement of Ofu soon after the loss of Lapita ceramics in Tonga. The question of whether the Lapita-PPW transition in Samoa represented two discrete settlement events, or a single transitional event, remained unresolved.

While new data using both ^14^C and new dating techniques (U/Th), combined with re-evaluation of existing data, provide great promise for resolving many chronological issues in the Pacific, the avoidance of key dating materials and the use of single-phase Bayesian evaluations unconstrained by stratigraphy or other form of independent dating control, effectively leave us with an imprecise chronology that is smeared over many decades. Ultimately, we are still left with an inability to truly resolve important chronological questions: 1. When was the earliest occupation? 2. How fast did people spread? 3. Was settlement continuous? 4. From what direction did settlement spread?

In this paper, we address some of these issues by presenting a re-investigation of the chronology of the To’aga site using a suite of new and precise shell and bone AMS ^14^C dates taken from key contexts. Based on these new ^14^C results, we evaluate the placement of To’aga within the current chronological model for the Late Lapita/Polynesian transition throughout Samoa and Tonga.

## The To’aga excavation and radiocarbon dates

The To’aga site (AS-13-1), situated on the southern coast of Ofu Island, was first test excavated in 1986 with more extensive excavations being undertaken in 1987 and 1989 [19]. The site consists of stratified cultural deposits situated within a coastal beach terrace located on the southern side of Ofu Island. The terrace was archaeologically investigated primarily through the excavation of 1-m^2^ test pits arrayed along a series of six transects running perpendicular to the coast (Fig 2). These transects revealed buried cultural deposits, some of which contained PPW ceramics, shell fishhooks, ornaments, and other artefacts and associated faunal remains. During the 1987 field season, an expanded trench was opened up to further sample the deeply buried, ceramic-bearing deposits, and is referred to below as the "Main Excavation". The original suite of radiocarbon dates from all three excavation seasons are presented in Kirch [20].

**Fig 2:**
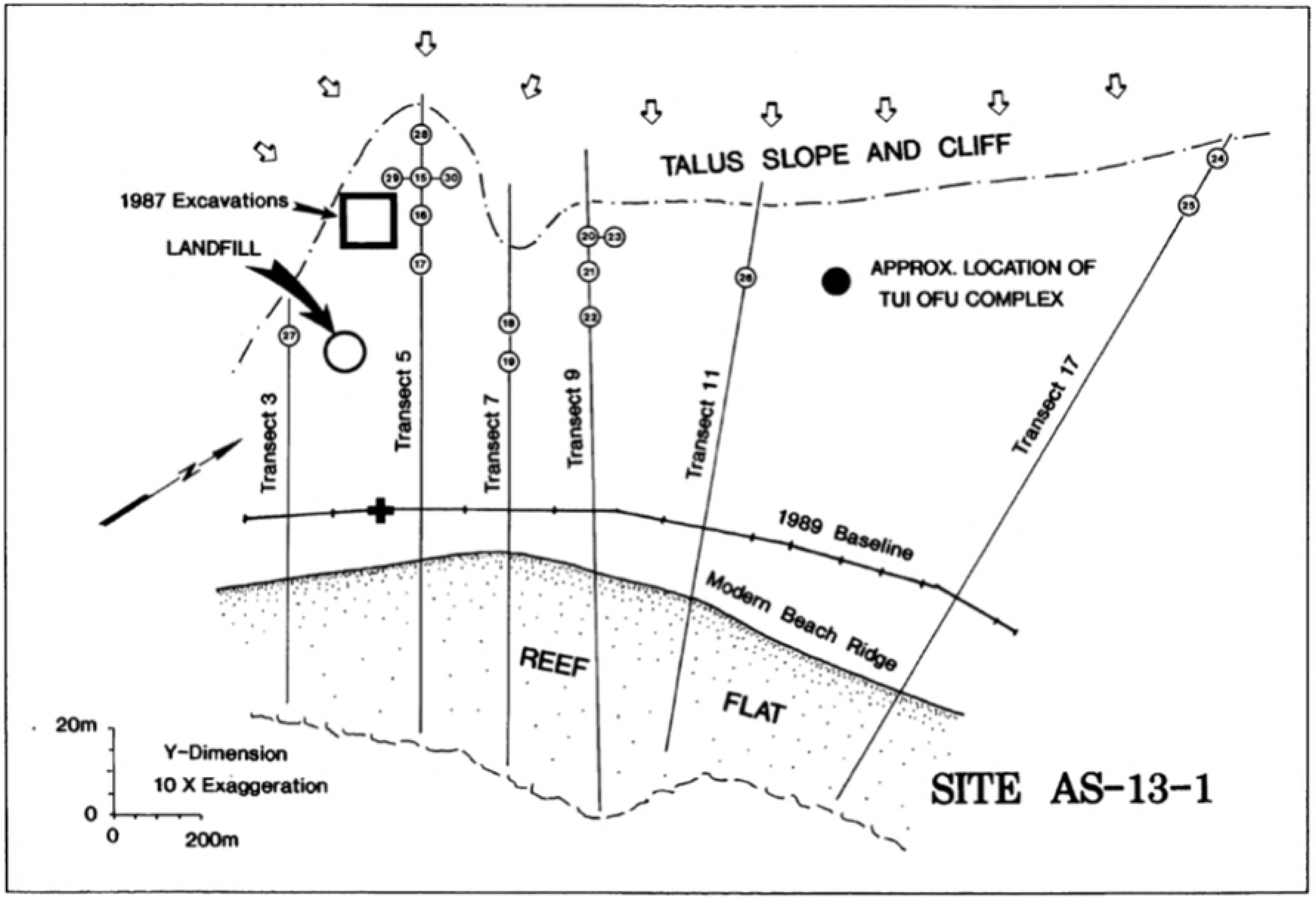
Map of the south eastern coast flat of Ofu Island showing location of excavations conducted in 1986, 1987, and 1989(from [25] fig 5.8)

For this renewed chronological evaluation, samples were sourced from excavated materials curated in the Oceanic Archaeology Laboratory at the University of California, Berkeley. These samples were specifically chosen to aid in defining the age of the oldest two main occupation deposits. They come from four locations; Unit 9 of the Main Excavation, Unit 10 located 45m to the southwest of the Main Excavation, Unit 28 located on Transect 5 to the east of the Main Excavation, and Unit 23 of Transect 9 also to the east of the Main Excavation (Figs 2 and 3). Radiocarbon dates and associated information is given in Table 1.

**Fig 3:**
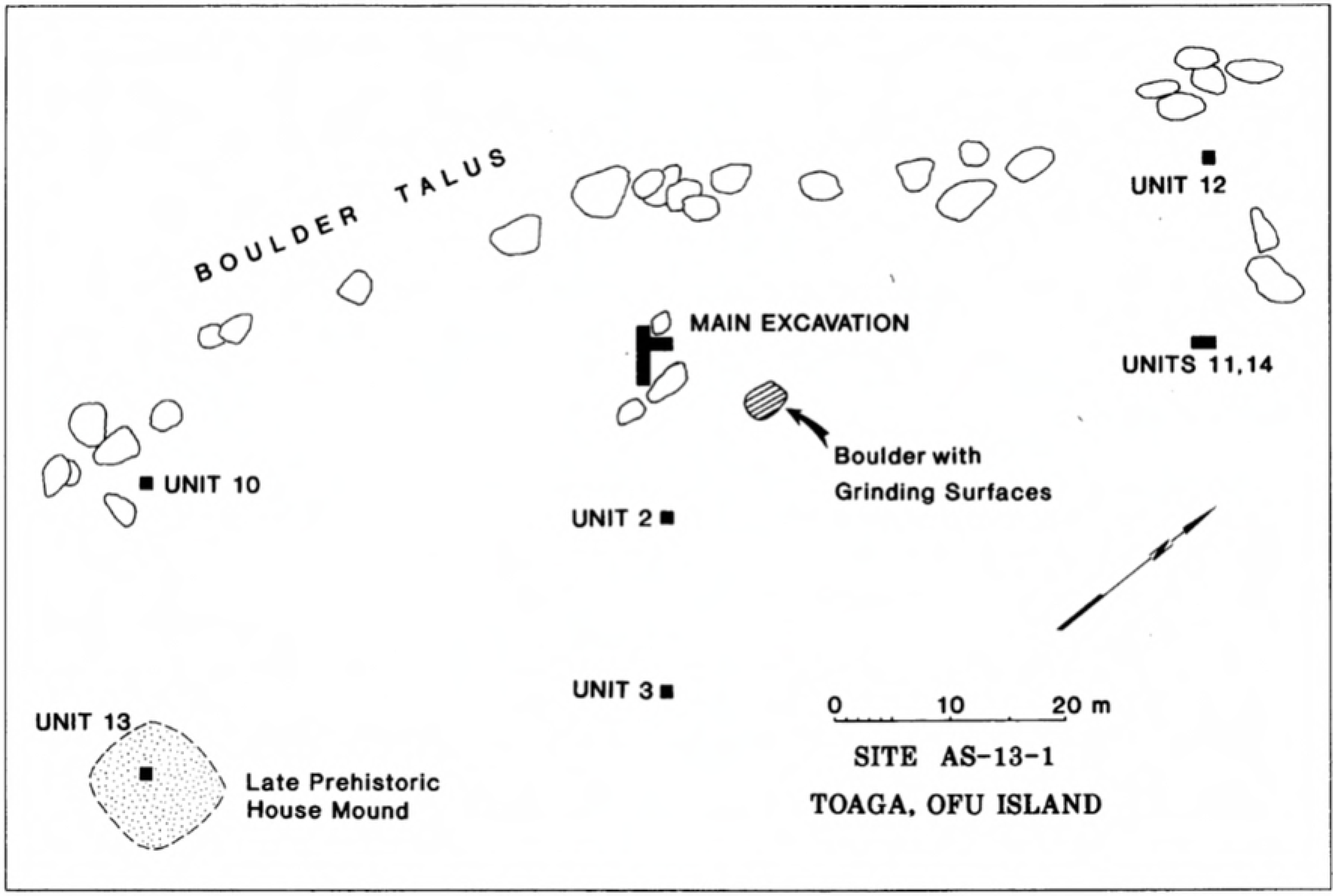
Map of the 1987 excavation area showing locations of Main Excavation and Unit 10 (from [25] fig 5.3)

**Table 1.** New and extant radiocarbon dates from earliest layers at To’aga, Ofu Island.

| Lab Code† | Provenance | Material | $\delta^{13}\text{C}$ (‰)* | $\delta^{18}\text{O}$ (‰)* | Conventional Radiocarbon Age (BP) | Unmodelled calibrated age 68% prob. (cal BP) | Unmodelled calibrated age 95% prob. (cal BP) |
| --- | --- | --- | --- | --- | --- | --- | --- |
| **1987 MAIN EXCAVATION** |  |  |  |  |  |  |  |
| Beta-25033 | Unit 6, Layer IIA-1 | Shell: *Turbo setosus* | 2.3 | - | 2640±80 | 2681-2433 | 2738-2322 |
| Beta-25034 | Unit 6, Layer IIB | Shell: *Turbo setosus* | 2.5 | - | 2570±80 | 2595-2333 | 2715-2210 |
| Wk-45468 | Unit 9, Layer IIB | Shell: Echinoid spine | 3.12 | -0.28 | 2720±16 | 2733-2596 | 2750-2480 |
| Wk-45469 | Unit 9, Layer IIB | Bone: *Rattus exulans* ** | -17.8 | - | 2344±16 | 2342-2184 | 2348-2159 |
| Beta-26464 | Unit 10, Layer IIB | Charcoal: Unidentified | -27.8 | - | 2620±140 | 2875-2489 | 3057-2351 |
| Wk-45472 | Unit 10, Layer IIB | Shell: Echinoid spine | 2.68 | -0.74 | 2761±16 | 2758-2655 | 2810-2525 |
| Wk-46708 | Unit 10, Layer IIB | Shell: *Turbo setosus* | 1.98 | -1.62 | 2576±16 | 2522-2345 | 2648-2326 |
| **TRANSECT 5** |  |  |  |  |  |  |  |
| Beta-35601 | Unit 28, Layer IIB (base) | Charcoal: Unidentified | -27.8 | - | 2900±110 | 3177-2881 | 3340-2788 |
| Wk-45473 | Unit 28, Layer IIB (base) | Shell: Echinoid spine | 2.25 | 0.34 | 2814±16 | 2720-2512 | 2764-2367 |
| Wk-46707 | Unit 28, Layer IIB (base) | Shell: *Turbo setosus* | 2.26 | -1.47 | 2819±16 | 2725-2520 | 2771-2373 |
| **TRANSECT 9** |  |  |  |  |  |  |  |
| Beta-35602 | Unit 23, Layer IIIA (Earth oven cut into B) | Charcoal: Unidentified | -26.9 | - | 2630±100 | 2866-2505 | 2958-2380 |
| Beta-35603 | Unit 23, Layer IIIB (base) | Charcoal: Unidentified | -28.4 | - | 2600±170 | 2869-2435 | 3156-2314 |
| Beta-35604 | Unit 23, Layer IIIB | Shell: *Tridacna maxima* | 1.7 | - | 2770±80 | 2703-2428 | 2794-2295 |
| Wk-45470 | Unit 23, Layer IIIB | Shell: Echinoid spine | 2.69 | -0.15 | 2809±17 | 2716-2508 | 2757-2365 |
| Wk-45471 | Unit 23, Layer IIIB | Bone: *Rattus exulans* | - | - | 2669±24 | 2780-2541 | 2844-2488 |
| Wk-47458 | Unit 23, Layer IIIC | Shell: Echinoid spine | 1.51 | -1.56 | 2892±25 | 2818-2612 | 2895-2460 |
| Wk-47459 | Unit 23, Layer IIIC | Shell: *Turbo* sp. | 1.99 | -2.08 | 2849±19 | 2749-2541 | 2833-2413 |

†Beta = Beta Analytic, Inc. (Beta-25673 and Beta-25035 are not included as results considered unreliable. The dating of *Asaphis violascens* [Beta-25035], a deposit feeder is also a possible cause of this erroneous age); Wk = Waikato Radiocarbon Dating Laboratory (New data).
\* Environmental $\delta^{13}\text{C}$ and $\delta^{18}\text{O}$ values reported with Wk- dates were measured on solid shell using a using a cavity ring-down $\text{CO}_2$ isotope analyser (CRDS) (Los Gatos Research model CCIA-46) at Waikato University using reference (NBS-19 and SDH synthetic $\text{CaCO}_3$ ). Measured precision of $\pm 0.35\text{‰}$ for $\delta^{13}\text{C}$ and $\pm 0.40\text{‰}$ for $\delta^{18}\text{O}$ . $\delta^{13}\text{C}$ and $\delta^{18}\text{O}$ values are reported as ‰ V-PDB.
\*\* $\delta^{13}\text{C}$ (VPDB) values for rat dietary correction were measured by Isotope Ratio Mass Spectrometry (IRMS) at Iso-trace Research Department of Chemistry, University of Otago on a Carlo Erba NA 1500 elemental analyser (EA), coupled with either a Europa Scientific '20/20 Hydra' or a Thermo Finnigan Delta Plus Advantage using reference (USGS-40, USGS-41) and control (EDTA-OAS and IAEA MP152) materials providing precision of $\sim \pm 0.2\text{‰}$ for $\delta^{13}\text{C}$ .

### Main Excavation

#### Original dates

The principal pottery-bearing layer in the Main Excavation is Layer IIB ([19] pg 51, fig. 5.5). This is capped by the culturally sterile Layer IIA, with a thin zone on top (IIA-1) that contained scattered shell and coarse-tempered sherds interpreted to be eroded material that accumulated following the abandonment of occupation in this area. Two dates on *Turbo setosus* shell from Layer IIB gave equivalent results (Beta-25033; 2311-2094 cal BP, and Beta-25034; 2244-2007 cal BP). Towards the base of Layer IIC large coral and volcanic rubble was interpreted as being the result of high energy deposition, most likely a storm event. Small numbers of thin, fine-tempered sherds were found in Layer IIC (a sandy, beach-ridge deposit) and in Layer III (massive colluvium derived from erosion up-slope); neither Layer IIC nor III were considered to represent in-situ occupations, and the sherds in them were interpreted as coming from the slope inland of the site. Layer IV was culturally sterile, and although two thin, fine-tempered sherds were recovered from Layer V these were also interpreted as having derived from an occupation locality further inland, now buried by colluvium and not accessible through hand excavation ([19] pg 51, 56). A date on a mixture of different shell taxa (Beta-25035; 3714-3549 cal BP) was obtained from lower Layer V, and a second date from nearby Unit 1 (Beta-25673; 3475-3326 cal BP) ([20] pg 91), but both results are considered to derive from secondary deposition. The Unit 10 excavation exposed a stratigraphic sequence very similar to that of the Main Excavation, with Layer IIB considered to be identical to Layer IIB in the Main Excavation units (dated by wood charcoal sample Beta-26464; 2916-2403 cal BP) ([20] pg 53-54).

#### New dates

An additional four samples from the main cultural deposits (Layer IIB) in Units 9 and 10 were selected for dating; two dates on echinoid spines (Wk-45468 and Wk-45472), one on *Rattus exulans* bone (Wk-45469), and one on *Turbo setosus* shell (Wk-46708).

### Transect 5, Unit 28

#### Original dates

Transect 5 lay approximately 100m east of the Main Excavation ([19] fig. 5.5). Unit 28 was placed at the base of a steep talus slope, and was specifically excavated to trace a deeply-buried cultural deposit exposed in Unit 15 (10m to the south of Unit 28) that contained a few sherds of thin, orange-slipped ceramics. Because the colluvial slope rises steeply along this inland edge of the coastal terrace, the excavation of Unit 28 required removal of 2.4m of colluvium and large boulders. Beneath the massive colluvium, the main cultural deposit (Layer II, with subcomponents IIB and IIC) contained ceramics and other artefacts and faunal material. Of 103 potsherds recovered from Layer IIC, 49 percent were fine, thin-wares. A single charcoal ^14^C sample from the base of Layer II (Beta-25601; 3257-2879 cal BP) was considered to date in-situ cultural material. Because of the steeply rising colluvial slope inland of Unit 28, further excavations to the north were not possible, but it was thought likely that older deposits remained unexcavated in that direction ([19] pg 60, 67).

#### New dates

Two dates, from the same context as Beta-25601 (Transect 5, Unit 28), were obtained to confirm the age of Layer IIB; a date on echinoid spine (Wk-45473) and a date on *Turbo setosus* shell (Wk-46707).

### Transect 9, Units 20/23

#### Original dates

As with Unit 28, Units 20/23 were also located at the base of the talus slope, but about 400 m east of the Main Excavation. Layer IIIA was associated with a large earth oven feature, dug down into Layer IIIB, containing large numbers of sea urchin spines (*Heterocentrotus mammillatus*), from which a charcoal ^14^C sample (Beta-35602; 2845-2612 cal BP) was originally dated. The underlying Layer IIIB was a thick, organic midden deposit that thinned into Layer IIIC ([19] pg 74, fig 5.22). Two charcoal dates (Beta-35603 and Beta-35602) from Unit 23, Layer III, returned ages of 2917-2382 cal BP and 2845-2612 cal BP respectively; significantly different from a shell date (Beta-35604; *Tridacna maxima*) from Layer IIIB (2444-2289 cal BP).

#### New dates

Four new dates were obtained from Unit 23; one on an echinoid spine (Wk-45470) and one on *Rattus exulans* bone (Wk-45471). Both came from the same context as the previous charcoal and *Tridacna* date (i.e., Layer IIB). Two additional dates, one on an echinoid spine (Wk-47458) and one on *Turbo* sp. shell (Wk-47459) from Layer IIIC were dated to provide a maximum age for this part of the site.

### Comment on Dates

#### Charcoal

Short-lived nut charcoal samples with only 1 year of growth are considered to be one of the most reliable dating materials, assuming there has been minimal stratigraphic displacement. It is also well-established that most wood charcoal determinations will date earlier than the event by an unknown amount [26]. This could be by a few years, or several hundred years, but Pacific research has indicated that, except in the highest precision analyses, minor inbuilt age typically goes unnoticed [27][28]. A large number of extant radiocarbon dates from Pacific contexts are on charcoal that has not been identified to short-lived materials, and removal of these unidentified charcoal dates from chronological evaluations would result in a dataset composed of relatively few dates. It has been demonstrated in Bayesian models that a small dataset has a greater detrimental impact on chronological resolution than the potential of unidentified charcoals to skew to older ages [29]. Outlier analysis specifically designed to account for this inbuilt age in charcoal is promising [5][29], but requires additional constraints (e.g., dates on short-lived materials or stratigraphy) against which to anchor the model (see also [30] where the model allows a small number of charcoal samples with inbuilt age to be younger than the context they represent, as would be the case with intrusive material).

#### Bone

*Rattus exulans* (Pacific rat) have a notoriety stemming from anomalously early dates from New Zealand archaeological contexts [31]. A range of theories have been put forward for these dates including the small size of these bones and potential laboratory contamination at the time of dating [32][33]. Twenty years later, there have been improvements in our understanding of dietary offsets and application of (dietary and reservoir) corrections to a range of animals that feed in both marine and terrestrial environments [34][9][35][36][37], as well as significant improvements in sample pretreatment and the abilities of AMS dating technology to date these tiny samples [38]. The importance of rats as human commensals, as evidenced by their presence in many archaeological deposits including To’aga ([19] pg 200, table 13.3), means there is significant benefit from being able to date these animals directly [39]. They do however, require a dietary correction which can be calculated from δ^15^N and δ^13^C analysis of the bone collagen and/or δ^13^C of bone carbonate [8]. We are not aware of any rat specific ^14^C studies into dietary corrections for island environments, but evaluation of ^14^C dates of other omnivorous animals (i.e., humans, pigs and chickens) from Pacific contexts [9][34] suggests a similar correction methodology is required.

#### Marine Shell

Marine shells that were gathered for food precisely date the timing of this activity. However, shell radiocarbon dates remain problematic because of uncertainties over ^14^C reservoir offsets (the ΔR value). This has resulted in many Pacific Island chronologies being dominated by wood and charcoal (e.g., [7][40][24][41][23]. See however, [5][21][42]). Petchey et al. [43] demonstrated little regional variation within the central Pacific Gyre zone, calculating an average ΔR value of 6±21 ^14^C years for modern shell collected from regions where currents are not interrupted by major island chains (e.g., Solomon Islands) or by contact between water bodies (e.g., Southland Front/Subtropical Front).

Subsequent research has demonstrated that shells, in particular, may give apparently erroneous ^14^C results depending on genera and even species selected, as well as local near shore conditions, unless appropriate corrections are applied. Within the South Pacific Gyre region one major concern when dating molluscs is the presence of ancient limestone on many islands [44], which can be taken up by animals either by directly living in waters that have percolated through the limestone, or directly by grazing on the rock [45][42]. Although *Turbo* and *Trochus* are potentially problematic in this respect ([46][47][48]) and sea urchins (echinoids) could similarly be affected due to their scavenging behaviour [49], the volcanic geology of Ofu Island negates any problems of this nature [50].

More recently it has become apparent that ΔR values have not remained stable over the last 3000 years; shifts in marine ^14^C between 3000- and 1900-years ago have been documented in corals from the eastern coastline of Australia that are linked to changes in ocean circulation [51][52]. These ΔR changes have also been documented in archaeological shell specimens from the central South Pacific Gyre region. Consequently, modern validation studies and reservoir correction values may not be applicable to archaeological material. Previously reported archaeological paired shell/charcoal ΔR values for Samoa (e.g., [53][20]) are excluded from this calculation because the paired charcoals selected for dating were not identified to short-lived materials. ΔR values obtained from U/Th dated corals from Ofu Island, reported in Clark et al. (2016), support these observations (Fig 4). The ΔR values available for the period between 3100 and 2650 BP have a pooled value of −47±10 ^14^C years (χ^2^_10:0.05_ = 33.33<18.31: GSD = 76). Between 2650 and 2250 BP the value drops to −161±11 ^14^C years (χ^2^_8:0.05_ = 13.29<15.51: GSD = 47). Unfortunately, this change occurs at a critical time for To’aga and the Lapita/PPW transition, and necessitates our testing both values.

**Fig 4.**
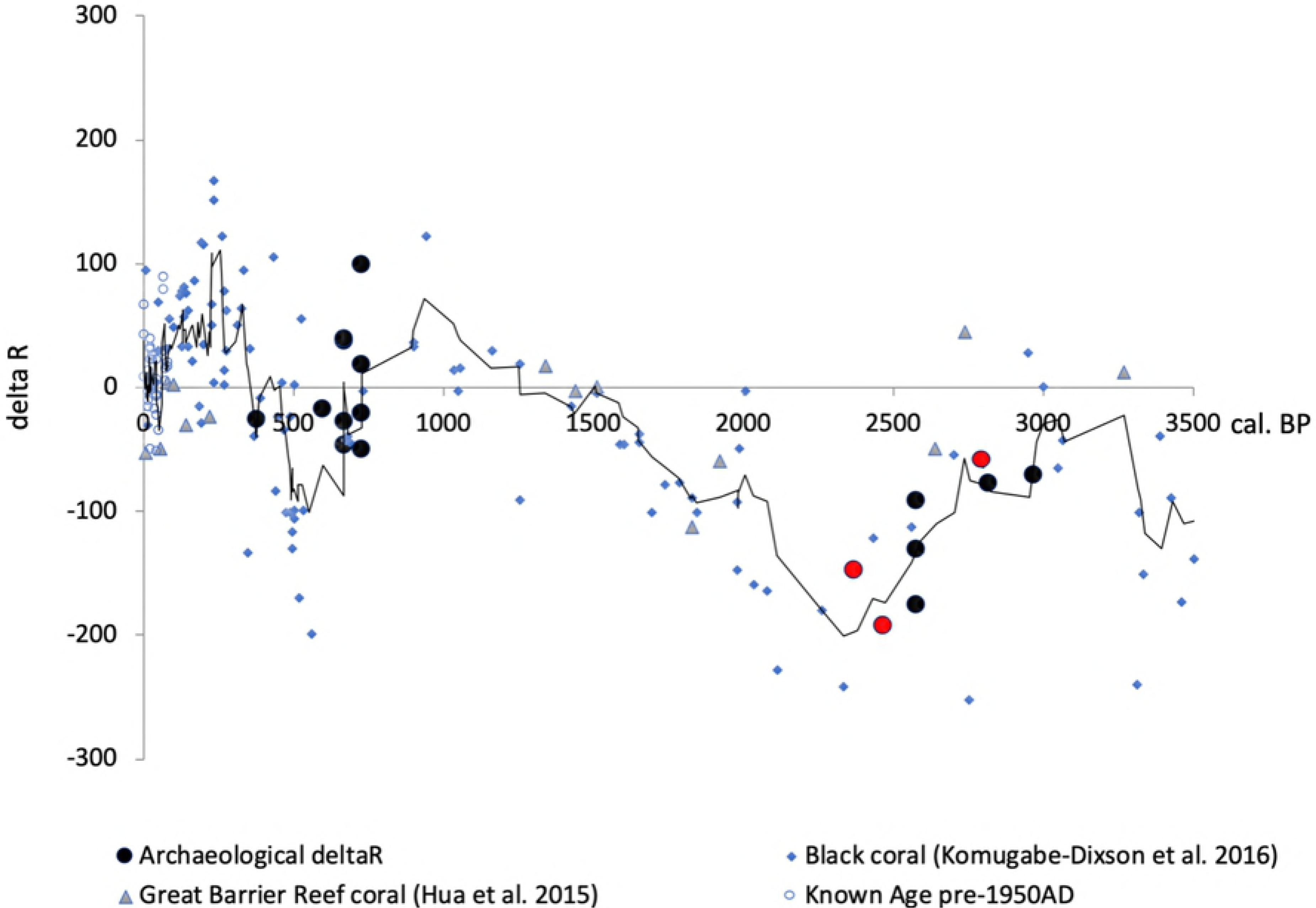
Change in delta R (ΔR) within the Pacific over the last 3500 years. Red circles = U/Th coral samples from Va’oto and Coconut Grove (Ofu Island). Black trendline based on a 4-point moving average.

#### Calibration

All radiocarbon dates were calibrated using OxCal v4.3.2 [54] with the Marine13 and Intcal13 curves [55]. We calibrated the shell results using either a ΔR of −47±76 ^14^C years or −161±47 ^14^C years. We have used the Northern Hemisphere ^14^C calibration curve (Intcal13) due to the position of Ofu within the Sub-Tropical Convergence Zone (after [56]), though a mixture of the Southern and Northern Hemisphere calibration curves is likely to be more appropriate (cf. [57]), but cannot currently be evaluated.

*Rattus exulans* bone dates require a dietary correction to the raw ^14^C data to obtain calibrated ages). A percent marine carbon (%MarineC, in this instance 28%) contribution to the diet was calculated for Wk-45469 which had a measured δ^13^C value of −17.8‰. A similar correction was assumed for Wk-45471 which was too small for both ^14^C and δ^13^C analysis. Following the methodology outlined in Petchey et al. [58] both dates were calibrated with a corresponding mixture of the Intcal13 and Marine13 ^14^C curves. The radiocarbon determinations, stable isotope values and calibrations, after the application of the resulting reservoir and dietary effects, can be found in Table 1. Dates are reported at 68% probability throughout the text.

#### Bayesian analysis

To provide the most probable chronology for To’aga, we conducted a Bayesian Sequence Analysis using OxCal 4.3.2 whereby radiocarbon ages are ordered on the basis of stratigraphic evidence [59]. In this model we have grouped the dates into two phases; Early (Transect 9, Unit 23 and Transect 5, Unit 28) and Main Excavation Layer (Units 6, 9 and 10) separated by a contiguous boundary. Within this “Early” phase, Unit 23 is further divided into three phases within a separate sequence that overlaps with the dates from Unit 28.

To assess the likelihood of any one sample being an outlier, a General t-type Outlier Model is inset into the sequence [60]. This enables outliers to be either too young or too old, and down-weights their influence in the model [26]. These dates are assigned a prior outlier probability of 0.05 and the scale of the offset is allowed to range anywhere between 10^0^ and 10^4^ years [“U(0,4)”]. The unidentified charcoal dates with possible inbuilt age are further assessed using an outlier correction for charcoal as described by Bronk Ramsey [26]; (Exp (1,–10,0), U(0,2.5),‘t’) whereby the exponential distribution runs from −10 to 0 with a time-constant of 1, ensuring outliers can only be older. The shifts are then scaled by a common scaling factor that can lie anywhere between 10^0^ and 10^2.5^ years. The impact of outliers on the model can be assessed by convergence values generated (S2-4). These should be >95%. Lower values indicate many different incompatible solutions to the model at these points. In Fig 5, calibrated ages before the model parameters have been applied (“prior probability values”) are shown as unfilled outlines. Posterior probability values after the model has been applied are shown in black.

## Results

There are differences in modelled age depending on whether a ΔR value of −47±76 or −161±47 ^14^C years has been used (Table 2 and S2 Tables). While these results do not conclusively support an age for the earliest deposits prior to 2650 cal BP, and therefore the suitability of using a ΔR of −47±76 ^14^C years, they do give strong support to this conclusion; use of the −161±47 ^14^C years ΔR value for the period 2650 - 2250 cal BP results in an even earlier date for the site (between 2817-2740 cal BP), which would then negate the use of this ΔR value. The slightly later deposits in Layer IIB in the Main Excavation are, however, likely to date to after 2650 cal BP, in which case a ΔR of −161±47 ^14^C years may be required for these layers. However, the Main Excavation Layer IIA-C deposits almost certainly include a mixture of early and later material based on the ceramic evidence (i.e., the proportions of thin-ware versus thick-ware). This caveat aside, we favour a third model where both ΔR values are applied depending on an early or late designation (Fig 5 and S 4). This produces a date for the initial settlement of To’aga of between 2782 and 2667 cal BP.

**Table 2:** Radiocarbon Bayesian model results for To’aga depending on ΔR used (68% prob.).

|  |  |  |
| --- | --- | --- |
| $A_{\text{model}} = 168.7$ | $A_{\text{model}} = 194.0$ | $A_{\text{model}} = 133.9$ |
| $A_{\text{overall}} = 131.6$ | $A_{\text{overall}} = 167.4$ | $A_{\text{overall}} = 96.4$ |
| No outliers | No outliers | 1 outlier (Wk-45469; *Rattus exulans*) |
| Boundary Start: **2764-2575** (67.6%) and<br>2566-2563 (0.6%) | Boundary Start: **2811-2740** cal BP | Boundary Start: **2782-2667** cal BP |
| Boundary Transition: **2627-2439** cal BP | Boundary Transition: **2754-2705** cal BP | Boundary Transition: **2727-2613** cal BP |
| Boundary End: **2301-2161** cal BP | Boundary End: **2338-2250** cal BP | Boundary End: **2329-2104** cal BP |
| $\Delta R = -47 \pm 76$ | $\Delta R = -161 \pm 47$ | Both |

**Fig 5:**
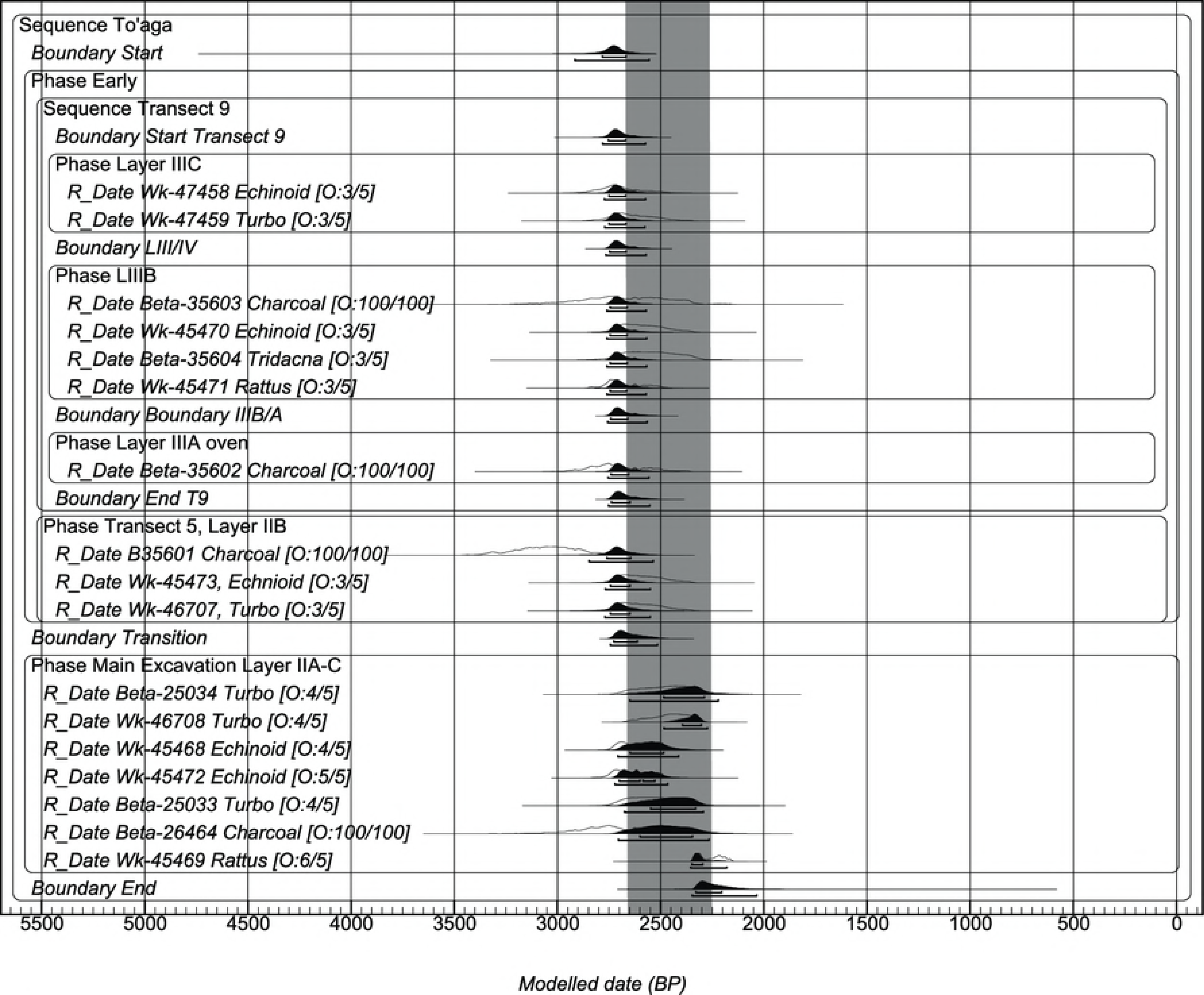
Bayesian sequence model for the To’aga site. Model uses a ΔR value of −161±47 ^14^C years for samples that date between 2650 and 2250 cal BP as indicated by the grey bar (i.e., “Main Excavation Layer IIA-C”) and −47±76 ^14^C years for earlier phases. 68% and 95% error margins are indicated by bars under each age distribution. The notation [O:2/5] indicates a 2% posterior probability of being an outlier in the model.

## Discussion

This multiphase sequence and outlier analysis provides the most secure age for the earliest deposits at To’aga available so far, and places first use of the site between 2782-2667 cal BP. This earliest phase of activity continues until 2613 cal BP (represented by boundary “Transition” in Table 2 and Fig 5). Subsequent activities in the Main Excavation take place between 2727 and 2104 cal BP (boundaries “Transition” and “End”), possibly incorporating material from many phases of activity that have become mixed together as a result of continual activity in the same general location. How do these new ages compare to the chronology of Lapita and PPW in Samoa and Tonga and is it possible to shed any further light on the key chronological questions for the region: when was the earliest occupation; how fast did people spread; was settlement continuous; and from what direction did settlement spread? In an attempt to answer these questions, and to highlight discrepancies in our current knowledge of regional chronology and radiocarbon methodology, the following regional comparison has been undertaken.

To ensure consistency across the datasets we have recalibrated the dates reported by Burley et al. [10] using the Intcal13 calibration curve. The original Tongan chronology presented by Burley et al. ([10][41]) uses the Southern Hemisphere terrestrial calibration curve [61]. Clark et al. [23] uses the Northern Hemisphere curve (Intcal13). The average difference between the two curves up to 1000 BP is 41±14 ^14^C years. We added the 16 new PPW dates reported by Burley et al. (2018), and while we have duplicated the overlapping phase Bayesian analysis used in the 2015 study we have also added outlier analysis to the OxCal code; specifying either General t-type Outlier for short-lived material or a Charcoal Outlier in situations where the charcoal has not positively been identified to short-lived material. Similarly, we have rerun the single-phase chronology based on short-lived materials presented in Clark et al. [62], again with outlier analysis, but excluding all To’aga dates. The Ofu PPW model includes relatively few dates (n=13) and is therefore at risk of being heavily biased by sampling and material choice. In particular, highly precise U/Th dates while providing a *terminus post quem* for the formation of the earliest cultural layers, are not definitively associated with cultural activity and are likely to have been deposited prior to first site use (For example, 2014-19 came from the boundary of the lowest cultural layer and the paleo beach ([62] pg 269). Consequently, in our revision of the Ofu PPW chronology we have only used coral that has been culturally modified. We have also applied a General t-type outlier to the U/Th dates to highlight that any date on this material dates the age of coral growth, not necessarily the age of the cultural modification (code for the Tongan Lapita, PPW and Samoan PPW sites is given in S 5). Lastly, the age of the Mulifanua turtle bone (NZA-4780; 3062 ± 66 BP) has been recalibrated using a ΔR of −47±76 ^14^C years.

Fig 6 shows several important differences between the old regional chronology (6a) and proposed new chronology (6b). The slightly increased age range for the three Tongan island groups during the Lapita phase is largely because of the large number of unidentified charcoal samples down-weighted by the outlier analysis. The slight shift to older ages, most obvious in the Tongan Lapita phase, is a consequence of using the Intcal13 curve. Now the earliest possible age for the Tongan PPW sites based on a single-phase analysis of all 45 Tongan PPW dates is 2744 cal BP. This refinement to the PPW sequence is the result of the increased number of dates now available and indicates the importance of larger numbers of dates when undertaking single-phase analyses of this type [29]. The duration of the Ofu PPW phase has been reduced by the removal of the culturally-unmodified coral U/Th dates (specifically 2014-19).

**Fig 6.**
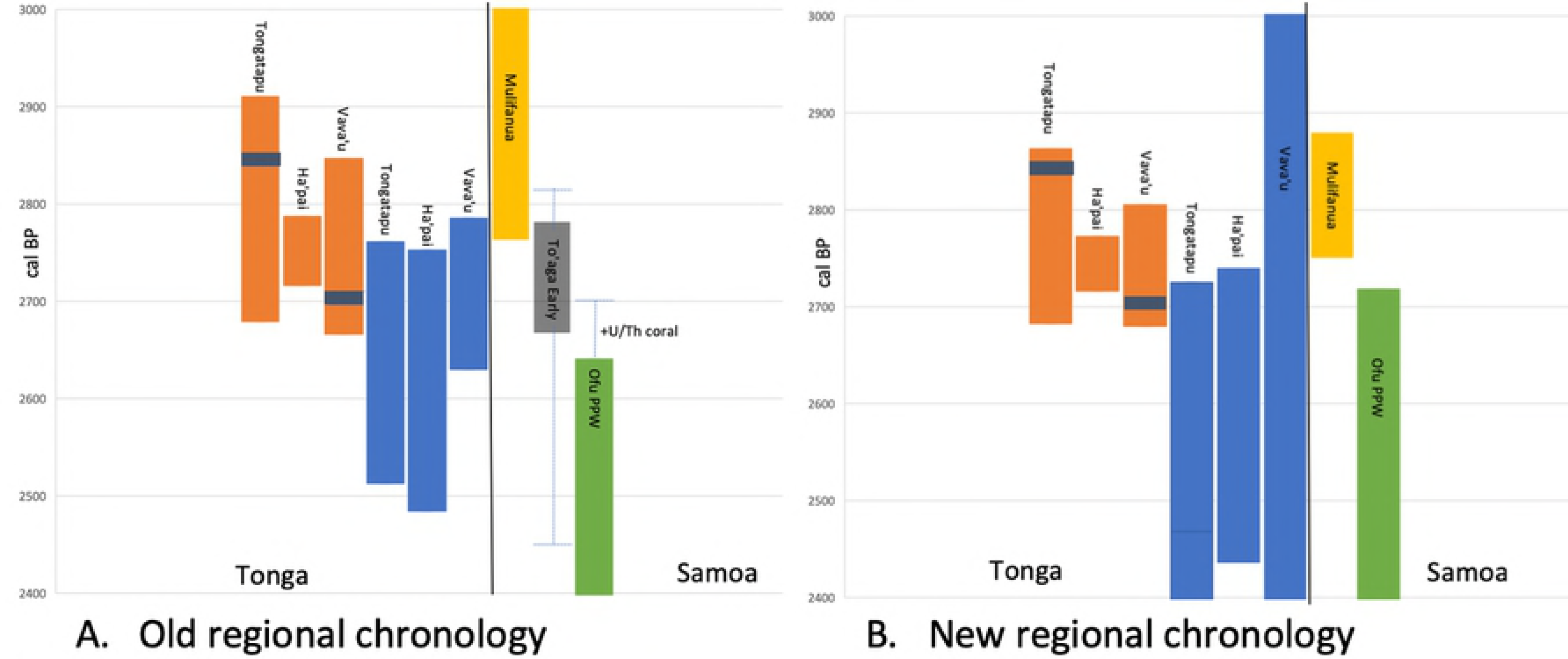
Regional chronology for Tonga and Samoa. A. Old regional chronology based on Clark et al. [62] for Samoan Polynesian Plainware sites (Va’oto and Coconut Grove), Mulifanua [16], and Burley et al. [24] for Tongan Lapita and PPW sites. B. New regional chronology using age appropriate ΔR for To’aga and Mulifanua (NZA-5800 only), Intcal13 for all charcoal samples, incorporating the data from Burley et al. [41] data into the PPW Tongan sites, and short-lived PPW dates from Ofu [62] excluding To’aga (dashed line shows extension of calibrated age if unmodified coral U/Th dates are included). The solid dark grey bar for To’aga represents the earliest phase of activity only. Dashed lines represent the maximum and minimum age range obtained using either −47 or −161 ΔR. Grey horizontal bars are U/Th dates considered by Burley et al. [24] to represent the most likely first Lapita landfall on Tongatapu and Vava’u.

The age for the Mulifanua Lapita site is now based on a single bone date rather than a pooled result, as calculated by Petchey (2001), and has shifted to a slightly older age range in response to the use of the −47±76 ^14^C year ΔR value as opposed to a ΔR of 57±23 ^14^C years used in Petchey (2001). Unfortunately, this date was produced over 20 years ago, and the available quality control information indicate that it would not pass the current bone assessment protocols essential for ^14^C dating (S 1). It should therefore be considered a minimum age.

### Directionality, Continuity and Speed

Burley et al. [24] argued for directionality with Lapita settlement moving from Tongatapu, where there was a short hiatus of ca. 70-90 years, before continued expansion through Ha’apai with near simultaneous movement into the Vava’u group; a maximum span of 158 years of Lapita settlement in Tonga. Our new model expands this to 243 years because of the additional uncertainty built into our outlier model. Regardless, U/Th dates from Tongatapu and Vava’u remain key indicators of the Lapita temporal limits. Lapita settlement of the remote northern islands of Niuatoputapu and Samoa were extensions of this movement ([24] pg 11). This argument implies initial Lapita colonization probably avoided northern Tonga and Samoa, but is difficult to reconcile given artefactual evidence of early connectivity between Fiji and Samoa [14][63]. Given the similarity in age of Mulifanua to early Lapita sites on Tongatapu our new model indicates that contact with ’Upolu most likely occurred early in the sequence, predating later Lapita settlements in Ha’pai and Vava’u, thus reopening the possibility of early movement via a northern corridor through Futuna and ‘Uvea [64][65]. This conclusion is tenuous, however, and necessitates further investigation of the antiquity of Mulifanua and these remote island outliers.

In the model presented by Burley et al. [24], the change to PPW occurred across the Tongan archipelago at a similar time. Our revised model is not currently precise enough to confirm that the transition to PPW occurred post-Lapita in Tonga, but it does support the conclusion of Burley et al. [24] that change was very rapid and near-simultaneous. The refined age for To’aga, and the likelihood that the current age range for the Lapita site at Mulifanua is a minimum age, suggests there may be a time lapse between these two early Samoan sites, but the exact magnitude is uncertain. Consequently, we cannot rule out abandonment of the Samoan islands for a short period before resettlement after a more secure base had been established in Tonga or elsewhere, but the gap identified by Rieth et al. [66] and Addison and Morrison [22] is now significantly reduced and an outright failure of Lapita colonisation of Samoa seems unlikely. Whether this possible “resettlement” at To’aga occurred at the terminal end of Tongan Lapita, or during the subsequent PPW phase is uncertain because the age for To’aga currently overlaps both. However, To’aga is significantly older than any other currently known Samoan plainware sites and fills a temporal gap between Lapita and PPW in Samoa. The continued use of To’aga from 2613 cal BP onwards (as evidenced by the Main Excavation/Transitional boundary date [Table 2]) clearly overlaps with activities elsewhere on Ofu Island. This continuum of human presence in Samoa has previously been postulated by Anderson and Clark ([67] pg 415) because the distinctive Samoan plainware ceramics would have necessitated some length of time to develop.

Even though we can refine the chronology movement through the islands using U/Th dating, as has been done for Tongan Lapita sites, this has not been possible for To’aga, and has not been as successful for defining PPW. In part, this is because of limited numbers of suitable culturally modified corals from key archaeological sites. Even the corpus of ^14^C dates on short-lived materials from early sites is limited (see [21]). While it is difficult to give any recommendations as to the number of dates required to improve our findings further, Schmid et al. ([29] pg 67) has suggested in such large-scale phase models ~280 dates will produce results of the highest precision, while the most accurate results are achieved where sampling density is uniformly distributed. Moreover, while many researchers question the usefulness of using ^14^C dating especially across the “Radiocarbon Plateau” (ca. 2650-2350 cal BP) there is structure present during this 300-year flat section of the calibration curve which could be utilised if larger numbers of precise (±20 years or better) dates on short-lived material were obtained from secure multiphase contexts. Clearly, more work is required across this region and time period.

## Conclusions

Our new dates and re-analysis of the site chronology indicates that the best estimation of the initial use of the To’aga site is between 2782 and 2667 cal BP. This confirms the antiquity of the site relative to other PPW sites on Ofu Island in Samoa, but we cannot confirm or refute an overlap in settlement timing with the Lapita site of Mulifanua on ’Upolu. Our findings also suggest that the initial occupation at To’aga was contemporary with the terminal Lapita/PPW transition in Tonga, but we cannot definitively say which came first. What is apparent, is that settlement in Samoa occurred early and is likely to have continued in a near unbroken sequence since Lapita times.

Over the decades, radiocarbon dates and the interpretation of those dates have become more sophisticated, but as research themes develop and new dating technologies become involved in the debates it has more than ever become necessary to refine the issues – both in archaeological research and radiocarbon methodology – that still plague the chronological interpretation. Chronometric hygiene methodologies initially provided the means to some clarity, but unfortunately removed a high proportion of dates from consideration. The limited number of early sites throughout the Pacific means we cannot afford to ignore the evidence already collected and where data are no longer considered of highest precision it is essential that extant excavations with curated samples are revisited. Bayesian methodologies now offer new opportunities to test these assumptions, as does refinement in ^14^C variation in both the marine and terrestrial reservoirs.

## Acknowledgements

We thank Margaret Norris, Rafter Radiocarbon Laboratory Manager, for providing data pertaining to the processing the Mulifanua bone date in 1994.

